# Impact of permagarden intervention on improving fruit and vegetable intake among vulnerable groups. A quasi-experimental study conducted in seven cities of Ethiopia

**DOI:** 10.1101/563965

**Authors:** Fikralem Alemu, Medhanit Mecha, Girmaye Medhin

## Abstract

**Background:** Increasing nutrient intake through home gardening is a sustainable strategy that can address multiple micronutrient deficiencies in developing countries. This study investigated the impact of permagarden intervention in increasing the frequency and diversity of vegetable and fruit consumption among vulnerable families.

**Method:** A quasi-experimental study was conducted from August 10 to September 30, 2015. A total of 884 care givers (427 from intervention and 457 control) participated in the study. Data was collected through face to face interview with caregivers of highly vulnerable children. Propensity score matching (PSM) was conducted using STATA software, and the program impact on the frequency and diversity of household’s vegetable consumption between intervention and control groups was assessed using chi square test.

**Result:** Intervention participants had a 13% greater increase frequency of one-week vegetables and fruits consumption compared with control participants (p<0.01). The diversity (consumption of 2 and more groups of vegetable and fruit) is higher among intervention groups than control groups (percentage difference=9, p-value<0.05). A significant higher percentage of participants in the intervention group reported getting the one-week vegetable and fruits mainly from their own garden (percentage difference 58.3, p<0.05). A significantly larger proportion of participants in the intervention group compared to the control group reported “high likelihood” on intention to grow vegetable in the future (percentage difference = 30%, and P<0.01). Perceived importance to include vegetables in every day meal is higher among intervention groups than control groups (percentage difference = 11.5%, P<0.01).

**Conclusions:** The observed higher frequency and diversity of household vegetable consumption among intervention group compared to control group suggests that nutrition and health programs need to promote household vegetable gardening as the means for address improve micro nutrient intake for vulnerable societies in least and middle developed countries.

## Introduction

Despite the growing body of evidence that suggests eating fruits and vegetables (FV) prevents several diseases, low consumption of fruit and vegetable remains one of the top 10 global risk factors for mortality (1, 2). The world health organization (WHO) defined low FV consumption of an individual as consuming less than the recommended intake of at least 400g of fruits and/or vegetables a day which is equivalent to five servings (3, 4). According to the WHO estimation, more than 2.7 million lives could be saved annually by increasing individual FV consumption (1). Undernutrition continues to account for 21% of the global disease burden among children in low and middle income countries (5). It is the major contributor of diarrheal morbidity among children less than five years of age, with a vicious cycle existing between diarrhea and under-nutrition (6, 7).

At a growing rate of 2.6% per annum and an estimated population of 104 million in 2018, Ethiopia is one of the largest least developed countries (LDCs) in Sub-Saharan Africa (70). About 10.2 million Ethiopians experienced food insecurity in 2016 (8). Ethiopia has more than 5 million orphans and vulnerable children (9) that could be prone to food insecurity (10). More than 51% of the death in under-five children attributed to undernutrition (11), and in 2016 an estimated 38% of children less than five years of age were stunted, 10% were wasted, and 24% were underweight (12).

Several studies revealed a number of positive health, social and recreational benefits of home gardening (13, 14). FV from home gardens have been documented as an important supplemental sources of nutrients, contribute to food and nutritional security, and allow families to produce their own fruits and vegetables organically (13, 15, 16). Vegetable gardens introduce more variety into the diets of children (17). Consequently, the intake of various nutrients such as vitamin A and calcium is increased with gardening (14). Home vegetable gardening has also psychosocial benefits. Learning gardening skills brings a sense of personal satisfaction to gardeners (18), and community gardens provide an opportunity to socialize with other community members beyond supplementing daily household consumption (19). Existing studies on home gardening were conducted in developed countries (20, 21). Limited evidence is available on the impact of home garden intervention in improving vegetable consumption of vulnerable groups in urban settings of low- and middle-income countries.

Yekokeb Berhan (2011-2017), was a USAID-program funded program that was designed to reduce the vulnerability of Highly Vulnerable Children (HVC) and their families in Ethiopia. Pact, the prime recipient of the grant, implements this program in partnership with FHI 360, Child Fund, and 49 local Civil Society Organization (CSOs) as well as with the Government of Ethiopia (22). A small scale permagarden intervention was implemented in response to the food and nutrition need of HVC and their families in 9 regions of Ethiopia where Yekokeb Berhan program was implemented (22). The current study is aimed to answer the questions “has permagarden intervention resulted in increased frequency and diversity of vegetable consumption?

## Material and Methods

### Study Design and setting

This is a post-intervention quasi-experimental study that involved intervention and control groups. Administratively, Ethiopia is divided into regions, zones, districts, and kebele where kebele is the lowest administrative unit (23). The study was conducted in 7 districts of two regions of Ethiopia, Amhara (5 districts) and Tigray (2 districts) where Yekokeb Berhan program has implemented. The national DHS report of 2016 showed that stunting, underweight, and wasting prevalence are higher in the two regions compared to the prevalence in other regions, and the national prevalence (24). The study included neighboring woreda as a control.

### Description of the Intervention

Caregivers of highly vulnerable children were trained on basic permagarden techniques for five days. The training topics covered different gardening aspects, namely, land preparation, garden layout, seedbed preparation, compost preparation, watering, weeding, and natural pest control. From April 2013 to May 2015, permagarden skill training was provided for 17,500 caregivers of HVC in the 9 regions of Ethiopia. Additionally, 817 trained volunteers delivered continuous consultation and follow-up to caregivers who begun gardening practice.

### Sample size estimation and sampling procedure

Sample size was calculated assuming an increase of 10 percentage points in the proportion of households exhibiting nutrition consumption as a result of the intervention and 50 percent of households in the control groups were assumed to consume FV. Using a standard parameter of 95% level of significance and 80%statistical power, the estimated sample size for intervention and control group was 388 each. An additional 10% was added to this sample size to cater for possible non-response and matching failures, making the total sample size 854 caregivers (427 from intervention and 427 controls).

Administratively, Ethiopia is structured into nine Regional Federal States and two City administrations. Each region is sub-divided into a set of zones, each zone is divided into set of woreda (an equivalent of a district) and each woreda is divided into kebeles. A *kebele* is the lowest government administrative structure. Two regions were selected randomly (Amhara and Tigray) from the total of six regions where Yekokeb Berhan program was implemented. We selected five *woreda* from Amhara region and two *woreda* from Tigray regions using simple random sampling technique. Proportional allocation of the study participants was done based on the number of caregivers trained and engaged in permagarden intervention per selected regions and woreda. Individual respondents were selected using systematic random sampling technique.

### Data collection methods and procedures

Data was collected through face to face interview using a structured questionnaire. The questionnaire included information about socio-demographic characteristics of participants; past and current engagement in home vegetable gardening practices, frequency of last one-week household vegetable consumption and the 24 hours FV consumption at household level. A questionnaire was originally developed in English and translated into the local languages (Amharic and Tigrigna). Supervisors and data collectors were provided with two days training about the study including field practice. The questionnaire was pre-tested with 50 households in 2 non-study neighborhood *woredas* that had similar context with the study area. The questions were assessed for clarity and suitability to the participants, and adjustments were made based on findings. During the data collection, completed questionnaires were evaluated by supervisors at the end of each day to ensure its completeness and consistency. Incomplete and inconsistent records were corrected in the field by re-visiting and re-examining study participants.

### Variable definition and measurements

#### Frequency of household FV consumption

The frequency of a one-week fruit and vegetable intake of the family was recorded as reported by caregivers with pre-coded responses (never, once, twice to three times, four to six times, one and at least once in a day). Their response was further recategorized as not frequent (1 to 3 times), and frequent (greater than 3 times).

#### Diversity of household’s FV consumption

Caregivers were asked to recall for themselves and their family if they had eaten specific FV in the last 24 hours according to FAO guideline (25). The groups of FV include group I (plant foods rich in vitamin A such as pumpkin, carrot, squash, or sweet potato, etc.), group II (plant foods rich in vitamin A fruits (ripe mango, cantaloupe, apricot, ripe papaya, etc.), group III (dark green leafy vegetables, including cassava leaves, kale, spinach, etc.) and group IV (any other vegetables e.g. tomato, onion, eggplant or other locally available vegetables). Households that reported consuming at least two FV groups were considered to have diversified FV consumption.

#### Future intention on home gardening

Future intention to be engaged in gardening practice is the likelihood that a household intends to practice vegetable gardening in the future. It was measured on a Likert scale of 1-5, where 5 stands for extremely likely and 1 stands for not likely at all (26). For reporting purposes response for intention were further recategorized in to less likely (scores 1 to 2), likely (score 3), and more likely (scores 4 to 5).

### Data management and data analysis

Cleaned data was double entered into Epdata software (EpiData Association, Odense, Denmark), and analyzed using STATA module psmatch2 developed by Leuven and Sianesi 2003 (27). Propensity score matching with nearest neighbor matching algorithm was conducted to balance the groups and minimize selection bias. Variables that were anticipated to influence outcomes including (a) ever trained on vegetable gardening by any organization, (b) household monthly income, (c) current participation in other gardening programs/intervention other than permagarden, (d) support from other source for vegetable gardening and participation in saving and lending groups were included in the questionnaire, and their influence was controlled during analysis The program impact on the frequency and diversity of household’s FV consumption between intervention and control groups was assessed using chi-square test and analysis. Percentage for categorical variables and mean values for continuous measurements were reported at 95% confidence level.

### Ethics statement

The research proposal was reviewed and approved by FHI 360’s Protection of Human Subjects Committee, and The National Ethics Review Committee (NRERC). Permission to conduct the study was obtained from the Amhara and Tigray regional women, children and youth affairs bureaus. Participants were informed about the purpose, their role in the study, the potential benefits, and possible risks associated with participating in this research study. A written informed consent was obtained from all participants for the interview and for the publication.

## Results

### Socio-demographic characteristics of participants

Socio-demographic characteristics of study participants is summarized in table 1. From the total of 884 households, 503 were from Amhara region and 381 were from Tigray region. The majority of the respondents (90%) were women, 40% from intervention and 50% from the control groups do not have formal education, 49.8% of the caregivers in the intervention group and 58% in the control group were not currently married (single/widowed or divorced).

**Table 1:** Socio-demographic characteristics study participants (caregivers) in Seven cities of Ethiopia, September 2015 (n=852).

| Households/Caregivers characteristics | Study group |  | Chi-Square test/ t-test | P-value |
| --- | --- | --- | --- | --- |
|  | Intervention (n=427) | Control (n=457) |  |  |
| **Region (%)** |  |  |  |  |
| Amhara | 55.3 | 58.4 | $\chi^2=0.95$ | 0.344 |
| Tigray | 46.7 | 41.6 |  |  |
| **Sex of caregiver (%)** |  |  |  |  |
| Male | 10.1 | 8.8 | $\chi^2=0.45$ | 0.502 |
| Female | 89.9 | 91.2 |  |  |
| Mean age of caregivers | 40.7 | 39.5 | t=1.51 |  |
| **Caregiver age group (%)** |  |  |  |  |
| Less than 30 | 12.0 | 15.4 | $\chi^2=2.58$ | 0.46 |
| 30-39 | 39.7 | 39.9 |  |  |
| 40-49 | 26.1 | 25.0 |  |  |
| 50 and above | 22.2 | 19.7 |  |  |
| **Caregiver relation to HVC (%)** |  |  |  |  |
| Mother | 71.6 | 79.1 | $\chi^2=8.3$ | P<0.05 |
| Father | 26.1 | 18.0 |  |  |
| Others | 2.3 | 2.9 |  |  |
| **Caregiver marital status (%)** |  |  |  |  |
| Married/Cohabit | 50.2 | 42.3 | $\chi^2=5.6$ | P<0.05 |
| Single/Divorce/Widow | 49.8 | 57.7 |  |  |
| **Caregiver level of Education (%)** |  |  |  |  |
| No formal Educ. | | | $\chi^2=20.8$ | P<0.001 |
| Elementary 1-8 | 40.0 | 50.6 |  |  |
| Secondary & above | 35.8 | 36.6 |  |  |
|  | 24.2 | 12.9 |  |  |
| **Caregiver occupation group (%)** |  |  |  |  |
| Daily labor | | | $\chi^2=12.1$ | P<0.05 |
| Gov. employee | 42.2 | 52.3 |  |  |
| Self-employed | 6.8 | 3.2 |  |  |
| Housewife | 27.4 | 21.7 |  |  |
| No occupation | 17.8 | 15.7 |  |  |
|  | 5.9 | 6.5 |  |  |
| Participate in saving group (%) | 69.8 | 48.5 | $\chi^2=41.3$ | $P<0.001$ |
| Run own small business | 37.6 | 19.0 | $\chi^2=36.2$ | $P<0.001$ |
| Household size (Mean) | 5.0 | 4.3 | $t= 5.15$ | $P<0.001$ |
| HHD monthly income in Birr (Mean) | 478 | 416 | $t=3.72$ | $P<0.001$ |

### Frequency of FV consumption

Figure.1 shows the frequency one-week FV consumption by intervention and control group. Out of the total 759 participants (404 intervention, 355 control), 88.4% of the intervention groups reported consuming FV at least twice in a week compared with 67.3% participants from control group (X2=49.6, P<0.01). 19% of households in intervention group consumed four and more times in a week compared with 6% in the control group (X2: 66.7, P<0.01). (Fig. 1)

**Figure 1.**
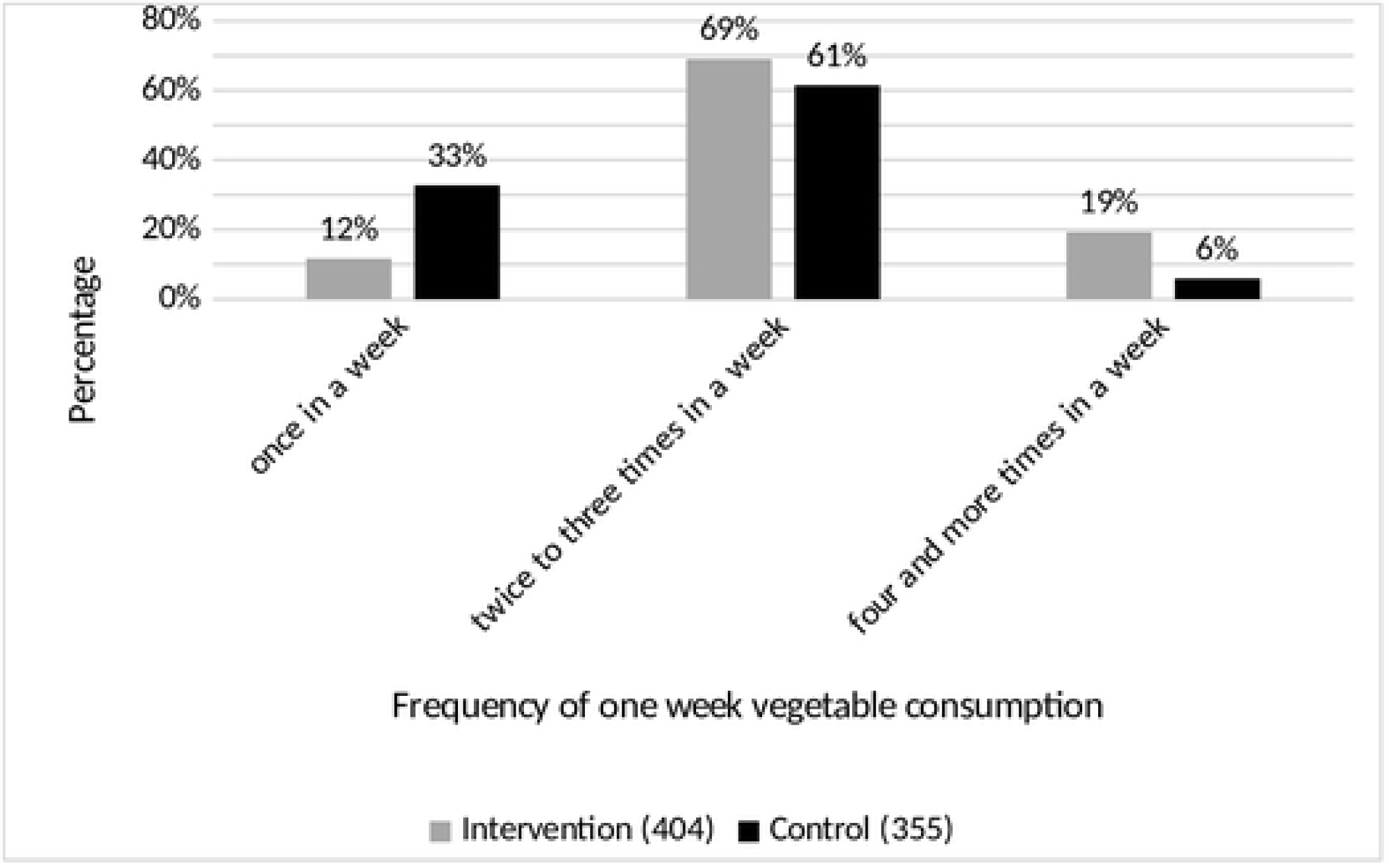
Frequency of vegetables and fruits consumption, September 2015

### Diversity of FV consumption

Figure 2 shows the 24 hours FV groups consumption by intervention and control groups. Results show that there was a significant difference between the intervention and control groups in the diversity of the 24 hours FV. Out of the total 626 participants (354 intervention, 272 control), fifty seven percent of participants in the intervention group reported consuming more than one group of vegetable (had a diversity of FV consumption) compared with fourty eight percent of participants in the control groups (x2= 5.304, p<0.05). (Fig. 2)

**Figure 2.**
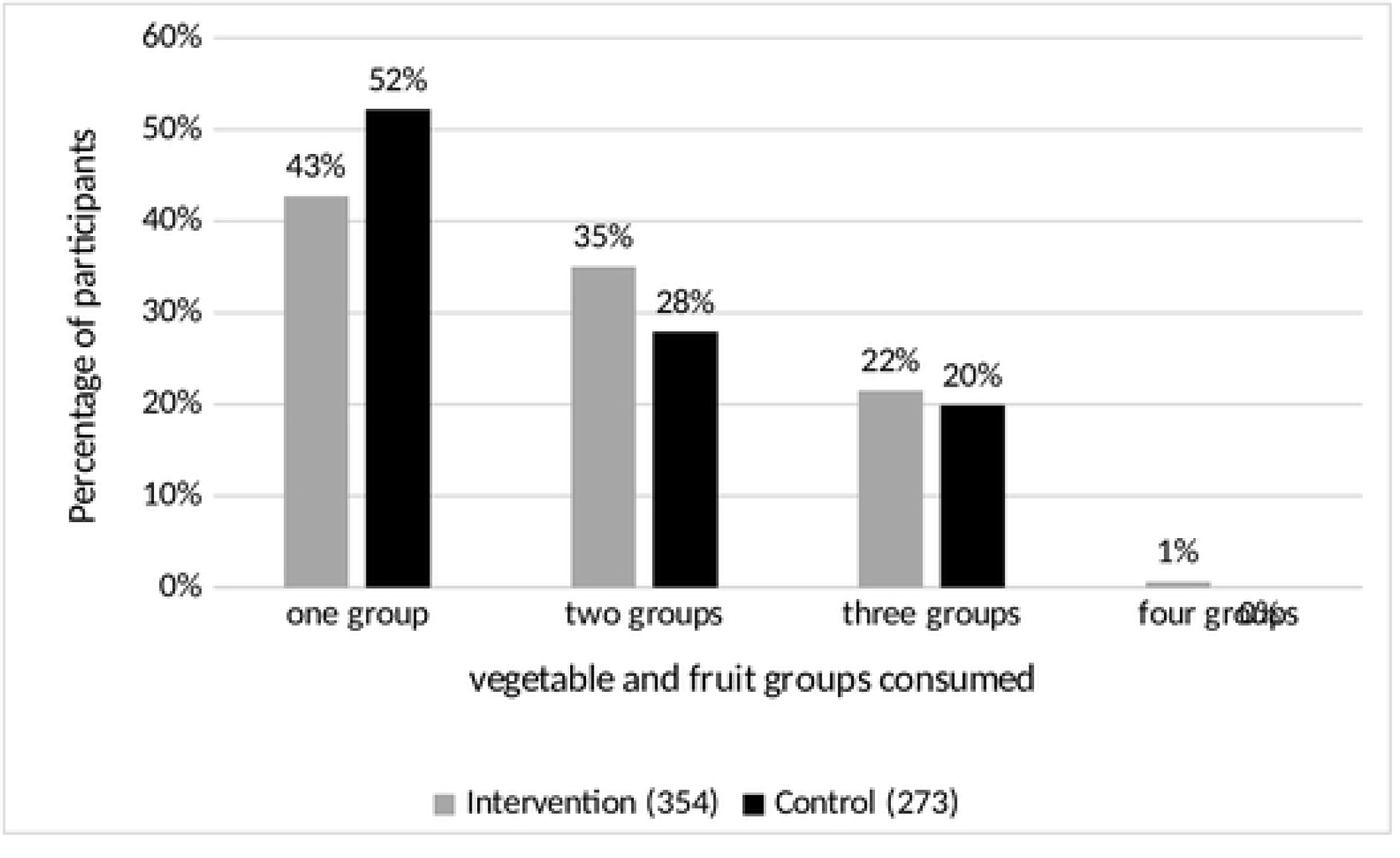
Diversity of 24 hours vegetable and fruit consumption among intervention and control groups, September 2015

Table 2 summarizes the specific FV consumed in 24 hours. There was no significant difference between the intervention and control groups in all the four FV groups consumed. The percentage of households that reported consuming vitamin A rich vegetable is almost equal to the percentage of HH in the intervention group. 98.3% of study participants in the intervention group have reported consuming dark green vegetables compared with 96.3% of HH in control group (P= 0.757). 42% of study participants in the intervention group have reported consuming Vitamin A rich fruits compared with 30.5% of HH in control group. (p< 0.01). 7.1% of study participants in the intervention group have reported consuming any other vegetables compared with 30.5% of HH in control group (p=0.754).

**Table 2.** Summary of the 24 hours household vegetable and fruit consumption by intervention and control groups, September 2015

| Fruits and vegetable dietary diversity | Intervention group | Control groups | Percentage difference | Chi-Square test | P-Value |
| --- | --- | --- | --- | --- | --- |
|  | Frequency/ % | Frequency/ % |  |  |  |
| Consumed at least 2 groups of fruit and vegetable. | 202(57) | 130(48) | 9 | 5.34 | $P < 0.05$ |
| Consumed vitamin A rich vegetables and tubers. | 113 (32) | 90 (33) | -1 | 0.096 | 0.757 |
| Consumed dark green leafy vegetables. | 348 (98.3) | 262 (96.3) | 2 | 2.425 | 0.119 |
| Consumed other vegetables | 25 (7.1) | 21 (7.7) | 0.6 | 0.098 | 0.754 |
| Consumed vitamin A rich fruits. | 149 (42) | 83 (30.5) | 11.5 | 8.836 | $P < 0.005$ |

### Access to FV from own garden

A significant higher proportion of participants from intervention groups (65.8%) compared with control groups (7.5%) reported getting vegetables and fruits from their own garden for their one-week consumption (x2=282, p<0.001). Consistently, the source of 24 hours vegetable and fruit consumption shows that fourty nine percent of participants in the intervention reported getting vitamin A rich plants from their garden compared with 6.7 percent participates in the control group (x2=43.5, p<0.001). Eighty two percent of intervention participants reported that they got the dark green vegetables from their garden compared with 12 percent of participates in the control group that reported getting the dark green vegetables from their garden (x2=291, p<0.001). Fifty percent of intervention participants reported that they got the Vitamin A rich fruits from their garden compared with 5 percent of participates in the control group that reported getting the Vitamin A rich fruits from their garden (x2=49.9, p<0.001). Twenty eight percent of intervention participants reported that they got any other vegetables from their garden compared with 0% participates in the control group that reported getting any other vegetables from their garden (x2=7.0, p<0.001). (figure 3).

**Figure 3.**
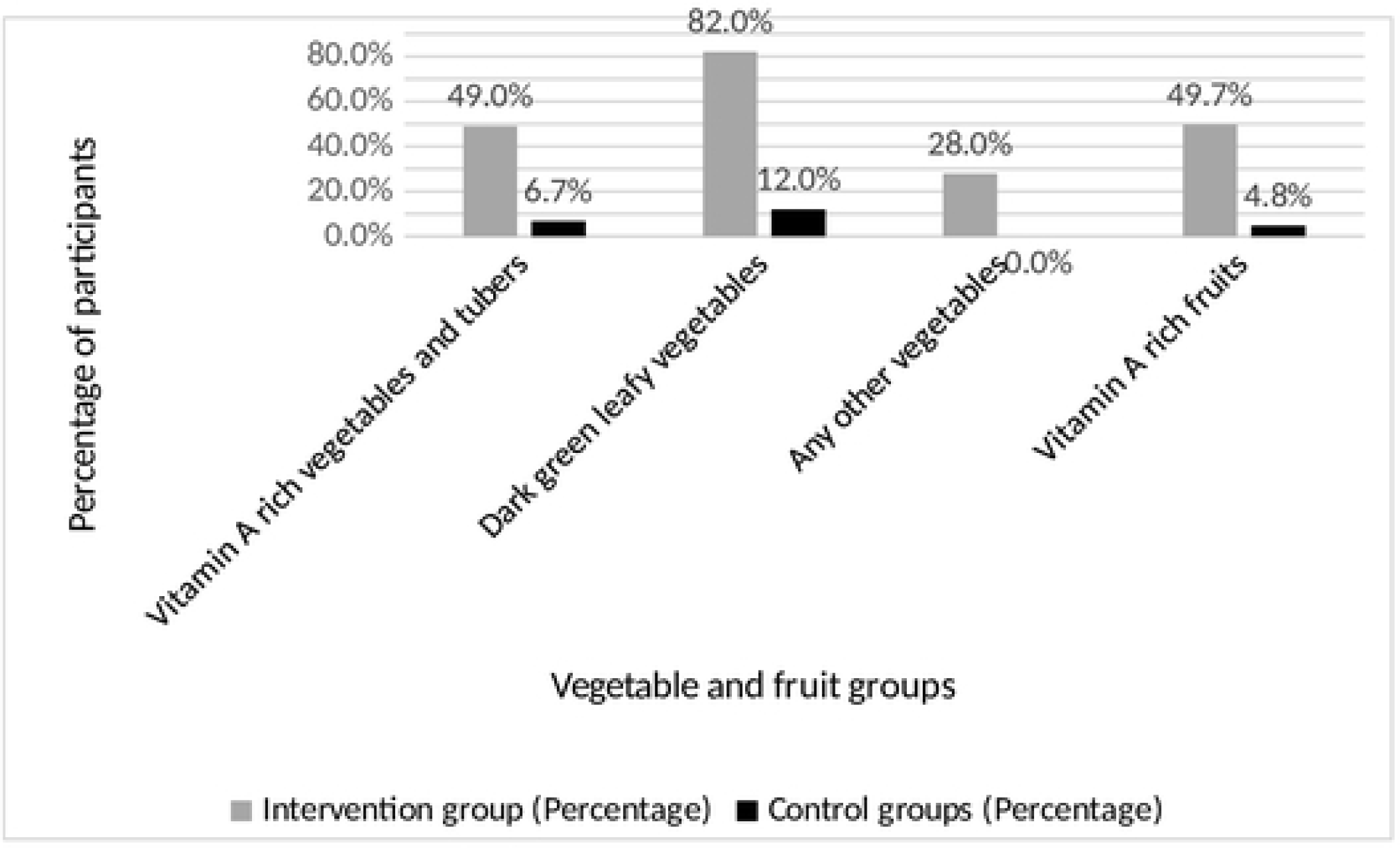
Percentage of participants with the source of vegetables and fruits and vegetables from own garden, September 2015

### Attitude on vegetable eating and future intention to home gardening

Table 3. shows perceived importance to include vegetables in every day meal and the likelihood of continue practicing home gardening in the future by intervention and control groups. Sixty six percent of participants in the intervention group and fifty four percent of participants in the control group reported “highly important” to include vegetable and fruit in every day diet (x2= 13.2, p<0.001). Regarding future intentions of participants on vegetable gardening, 58% of the control group and 88% of the intervention group reported high likely’ to continue home gardening in the future (x2=89.7, p<0.001).

**Table 3.**
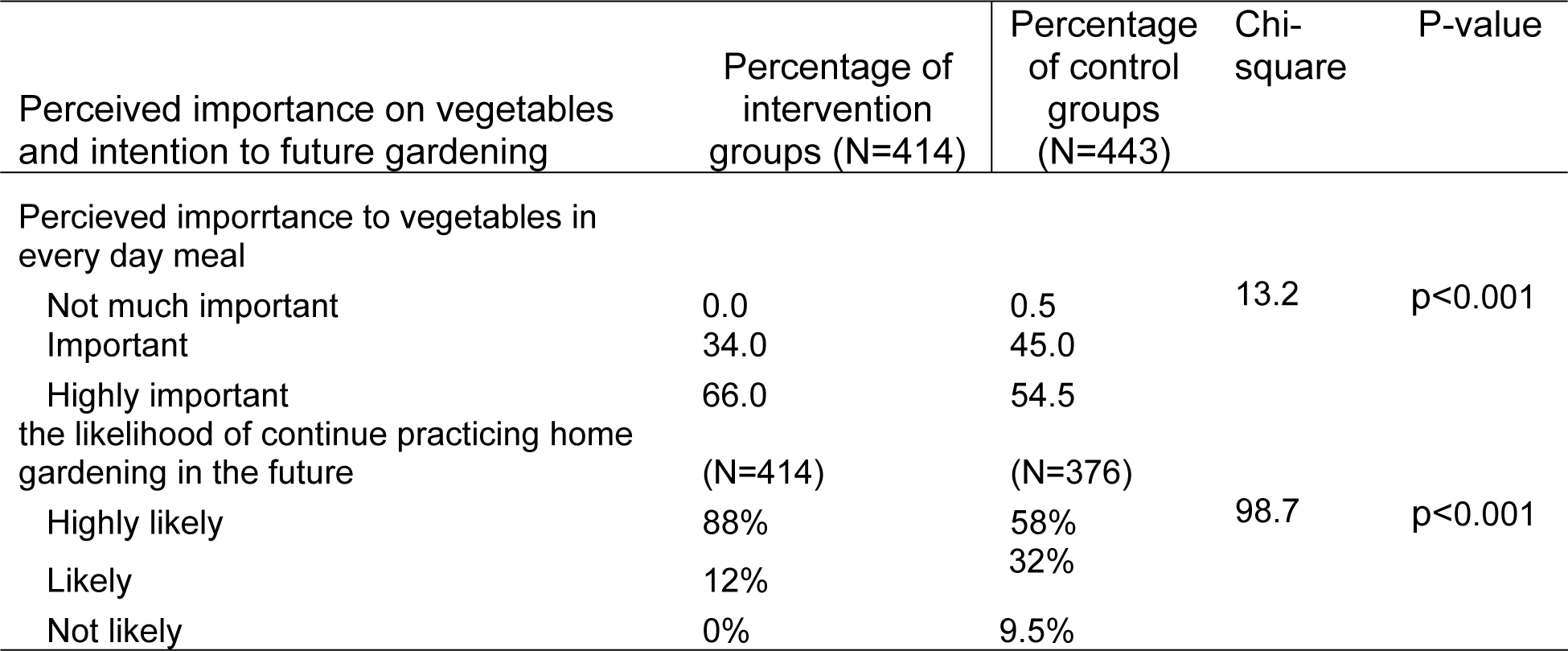
Perceived importance to include vegetables in every day meal and the likelihood of continue practicing home gardening in the future by intervention and control groups, September 2015.

| Perceived importance on vegetables and intention to future gardening | Percentage of intervention groups (N=414) | Percentage of control groups (N=443) | Chi-square | P-value |
| --- | --- | --- | --- | --- |
| Perceived importance to vegetables in every day meal |  |  |  |  |
| Not much important | 0.0 | 0.5 | 13.2 | $p<0.001$ |
| Important | 34.0 | 45.0 |  |  |
| Highly important | 66.0 | 54.5 |  |  |
| the likelihood of continue practicing home gardening in the future | (N=414) | (N=376) |  |  |
| Highly likely | 88% | 58% | 98.7 | p<0.001 |
| Likely | 12% | 32% |  |  |
| Not likely | 0% | 9.5% |  |  |

## Discussion

This study was conducted among caregivers of highly vulnerable children in seven *woreda* of Ethiopia and examined whether the groups that implement permagarden have higher FV intake compared with their control. Results show that intervention groups have higher frequency and diversity of fruit and vegetables intake and higher intention to practice gardening in future.

The study found that permagarden intervention groups ate FV more frequent times per week than control groups. A significant higher percentage of participants in the intervention group reported getting their family consumption of FV form their own garden. Consistent to the current finding, other researchers have shown that the most significant impact of home gardening was its impact to access to fresh produce FV (17, 28). The current finding, and other research suggests that home gardening can contribute to improving food security by helping to provide access to variety of FV to vulnerable households and their families (29–31).

There was a significant difference between intervention group and control group in the diversification of FV consumption (consumed 2 and more group). However, the diversification is still low in both groups. out of the total participants, 42% of intervention and 52% of the control groups consumed only one type of vegetable, and 35% of intervention and 28% controls consumed only two groups of FV groups. The possible reason for this finding might be lack of knowledge on the nutrition values of different groups of FV. This conclusion can be strengthened by findings of other research that reported home-garden interventions are most effective when combined with promotional and educational interventions (32). Based on the result from this study, and other studies, we suggest that home garden programs should integrate nutrition education (13, 33).

This study shows that the participants intention to continue home gardening and perceived importance to include vegetable in every day meals were significantly higher among intervention groups compared to the control group. Participation in the permagarden intervention might have influenced participants to develop positive attitude on FV consumption. Consistently, other studies reported that that participating in gardens increases preferences for vegetables (34, 35).

Several research have shown that the poorest people had the highest prevalence of low FV consumption (36–38).This study provides evidence that home gardens may have potential as a nutrition intervention to increase FV intake among the poor. The permagarden intervention was implemented integrated into Orphan and Vulnerable Children (OVC) program that has aimed to increase access to food and nutrition. Thus, making the model viable in urban context to address the food and nutrition gap among vulnerable societies of least developed countries (39). Since the current findings are based on post intervention study using a one-time data, evidence from future randomized intervention study with baseline data are likely to generate stronger conclusion with programmatic implications.

### Strength and limitation

A strength of the current pilot study was its ability to control confounding factors through PSM. Several study limitations may affect the interpretation of these results. The study was not a randomized controlled trial. Based on a one-time post-intervention data in this study, it might be difficult to establish causal relationship between the exposure and observed outcomes. We have estimated vegetable consumption based on the participant’s self-reported responses that might reduce the level of confidence we can put on the strength of intervention effect.

### Conclusions

The observed higher frequency of household FV consumption and increased access to family consumption FV form own garden among permagarden intervention group compared to control group suggest that nutrition and health programs need to promote household vegetable gardening as the means for address improve micro nutrition intake of vulnerable societies in least and middle developed countries.

## ACRONYMS

CC: Community Committee
CCC: Community Care coalition
CF: Community Facilitators
CSI: Child Support Index
CSO: Civil Society Organization
FHI 360: Family Health International 360
FV: Fruits and Vegetables
GOE: Government of Ethiopia
IEC: Independent or Institutional Ethics Committee
HVC: Highly Vulnerable Children
IRB: Institutional Review Board
NIH: National Institutes of Health
PHSC: Protection of Human Subjects Committee
OVC: Orphan and Vulnerable Children
SD: Standard Deviation
TOT: Training of Trainers
UNICEF: United Nations Children’s Fund
USD: US Dollar
USAID: United States Agency for international Development
WHO: World Health Organization

## Declarations

### Ethics approval, consent for participation and publication

The research proposal was reviewed and approved by FHI 360’s Protection of Human Subjects Committee, and The National Ethics Review Committee (NRERC). Permission to conduct the study was obtained from the Amhara and Tigray regional women children and youth affairs bureaus A written informed consent was obtained from all participants for the interview and for the publication.

### Availability of data and materials

All the necessary data were analyzed and included in this manuscript. For further need, data can be obtained from the primary author upon request.

### Authors’ Contributions

FA led the study design, analysis and write up of the manuscript. MM and GM participated in the review and guidance of the design, analysis and manuscript write-up. All authors read and approved the final manuscript.

## Acknowledgement

We would like to acknowledge Mr. Peter Jensen for providing permagarden training and follow up of most of the training site. This study wouldn’t be practical without volunteer participation of study participants and data collectors. We also like to acknowledge the contribution of Yekokeb Berhan regional staffs and implementing partner staffs for their involvement and support during data collection.

